# Flock: A Negative-Enriched Protein–Protein Interaction Dataset

**DOI:** 10.64898/2026.09.30.755672

**Authors:** Naail Kashif-Khan, Holly Gregory, Andrew Potterton

## Abstract

Protein–protein interactions (PPIs) govern fundamental biological processes and are central to therapeutic discovery, yet computational prediction methods remain severely limited by a lack of high-quality negative data (non-interacting PPIs). Existing models are trained almost exclusively on positive interactions or random pairs designated as negatives, while legacy negative datasets are limited in scale and contain correctness errors. To address this, we introduce Flock, a negative-enriched PPI dataset of unprecedented scale, containing 26,934 positive and 377,643 negative pairs. Flock integrates biological assemblies from the PDB with highly confident, curated negatives mined from scientific literature using a state-of-the-art agentic LLM workflow. To address the evaluation leakage seen in protein structure modelling, we also present Flock’s Leakage-Free Set, a benchmark that strictly controls for sequence and interface similarity, and historical date-based cutoffs used in the training of co-folding models. Our evaluation reveals that the co-folding model ESMFold2 shows performance degradation on the Leakage-Free Set, whereas the protein language model ESMC demonstrates stronger generalisation. Flock provides the most comprehensive and challenging PPI benchmark to date, establishing a rigorous new standard for evaluating the generalisability of protein structure and language models on this task.

## 1 Introduction

Protein–protein interactions (PPIs) govern how biology works, dictating cell signalling pathways, gene regulation, protein function and enzymatic activity. They are also important in drug discovery, particularly in molecular glue therapeutics. Molecular glues often strengthen weak, endogenous PPIs, leading to protein degradation, reactivation or cell death [1]. A better understanding of PPIs could aid in the discovery of novel molecular glue therapeutics.

Experimental PPI determination is a challenging task, due to the large number of estimated PPIs, the time and cost required to generate data, the scarcity of non-interacting protein data [2] and the sometimes inconclusive nature of experimentally-derived data [3]. To this end, there have been a variety of computational models developed for predicting PPIs, including co-folding [4–6] and protein language models [6]. However, the majority of PPI prediction models are trained on positive data only due to the aforementioned paucity of negative data [2]. Instead of using explicit negative pairs, those models [4–6] use random pairs which by definition may include false negatives.

Another common issue in training regimes for PPI models is lack of rigorous training and validation splits [7]. It was revealed that simple features such as cellular localisation could predict PPIs as accurately as a trained model [8]. Similarly, co-folding models are trained using a date-based cutoff rather than any consideration of a held out protein sequence or structure set [4, 5].

**Contributions**. This paper provides the following contributions:

1. A dataset called Flock that collates known PPI positives (from PINDER [9] and PPI3D [10]) and negatives (from the PDB and mined agentically from scientific literature). This marks the first negative PPI data update since 2013 and the debut of LLM agents for literature mining of those negatives.
2. A robust benchmark (Flock’s Leakage-Free Set) for the PPI prediction task that considers both interface similarity and training date cutoffs from co-folding models.
3. Evaluation of the performance of a co-folding model and ESMC [6] embeddings on Flock’s Leakage-Free Set and comparison against a leaked set.

## 2 Related Work

### 2.1 Non-Interacting Proteins (Negative PPI) Dataset

Both the original Negatome [11] and Negatome v2 [2] datasets used curated literature and PDB data to define non-interacting proteins. Natural language processing was used to mine scientific literature to search for negatives, an approach known to be greatly surpassed in accuracy by LLMs [12]. Non-interacting protein pairs were identified from non-interacting chains in PDB biological assemblies. Due to the substantial increase in PDB structures since 2013, aided by Cryo-EM determination of large complexes, the yield of non-interacting proteins is expected to far exceed Negatome v2.

### 2.2 Positive PPI Datasets

PINDER is a large dataset of structures of positive PPIs derived from PDB assemblies and augmented structures [9]. Interface similarity was used to create a PINDER-XL benchmark without data leakage. The dataset is mainly designed for protein–protein structure prediction.

PPI3D [10] is a regularly updating database designed for searching, analysing and modelling protein–protein, peptide, and nucleic acid interactions using structural data taken from the PDB and SCOPe [13].

PRING compiled and filtered databases including UniProt, STRING, IntAct and Reactome into a network of interactions rather than a binary classification task [14].

## 3 Methods

### 3.1 Derivation of negatives from the PDB

Following the methodologies of Smialowski et al. [11] and Blohm et al. [2], non-interacting pairs derived from the PDB were regenerated. In contrast to the published chain-level procedure, these pairs were aggregated at the protein level. A pair was emitted exclusively when every backing chain pair was non-interacting, which prevents a pair from being designated as negative while another copy of the same two proteins is in contact.

### 3.2 Derivation of negatives from scientific literature using LLMs

A starting corpus of papers was compiled by scraping Europe PMC using a set of search strings (Appendices B and C.1), each with a matched regular expression (regex). Regexes were used to recover the sentences matched by the search strings in each paper, and the paragraph containing them. In total, 365,743 text blocks from 150,217 papers were retrieved for further mining. Claude Sonnet 5 was used to extract potential non-interacting pairs from each sentence and its surrounding paragraph, resulting in 147,905 potential negative PPI pairs (Appendix B.1).

These pairs were first deterministically filtered (Appendix B.2). 10,152 pairs were deterministically assigned a pair of UniProt accessions. Of the remainder, around 30,000 pairs had at least one ambiguous member where the text name only resolved to a list of possible UniProt accessions. In total, 38,766 individual protein names were resolved to UniProt accessions using an LLM screen (Appendix B.1). This set of 27,077 pairs from 14,947 papers was subjected to a final agentic LLM curation pass (Appendix B.1). This resulted in a final curated dataset of 10,480 usable negative literature pairs. Appendix C outlines the experiments carried out to decide the LLM models and settings used in screens and curation.

### 3.3 Gathering of positive PPIs from PINDER and PPI3D

The union of PINDER [9] and PPI3D [10] data were used as sources of positive PPIs. Only biologically relevant interfaces were selected, and crystallographic artefacts were removed. This is prelabelled in PINDER, and was calculated for PPI3D using PRODIGY-crystal 1.0.1 [15]. The following filters were applied to protein pairs: resolution ≤ 4.5 Å, ≥ 4 intermolecular contacts, buried SASA ≥ 200 Å^2^ or Voronoi area 100 ≥ Å^2^, mappable to UniProt accession code and not an expression tag (GFP, GST, FRP, MBP, SUMO-1/2/3, Smt3). Only unique pairs passing the above criteria were included in Flock.

### 3.4 Creation of Flock, Leaked and Leakage-Free Sets

Negatives from the PDB and literature were first filtered to remove any pairs found as positives in IntAct or Flock’s own positives. Homodimer pairs and duplicate pairs were also removed.

From this Flock set, we then generate a Leakage-Free subset. The latest co-folding training cutoff date of Boltz-2 (2023-06-01) [16] was used to filter both positive and negative pairs. Positive pairs were removed if they were similar to a pre-cutoff entry, with similarity defined by membership of the same interface cluster in either source. Clustering was applied in the following way: PPI3D, subunits clustered at 40% sequence identity and 50% interface contact similarity; PINDER, chain pairs grouped by Foldseek structural clusters (mean lDDT ≥ 0.7 with ≥ 50% interface residue overlap). Negatives were removed if a pre-cutoff entry included both proteins or contained two chains mapped to different accessions homologous to both sides (*>*40% sequence identity at ≥ 50% coverage). Finally, we filter the Leakage-Free set to a list of targets, such that each target has at least 2 positive partners and at least 20 negative partners.

To create an equivalent leaked set for comparison, only pairs that are in the training set of Boltz-2 (2023-06-01) or have a homolog in the training set are included. The same ratio of positive and negative pairs were used as a filter.

### 3.5 Validation of existing models on Flock’s Leakage-Free Set

ESMFold2 was used as a representative state-of-the-art co-folding model [6]. Fast-mode with a single sequence (`biohub/ESMFold2-Fast` model weights, no MSA) was used. The following settings were used: 3 recycling iterations, 50 diffusion steps, 1 number of diffusion cycles. Recalculated ipSAE values were used as a scoring metric based on chain length (`iptm_d0chn`) [17].

ESMC’s PPI prediction methodology (Section A.4.4.3 of Candido et al. [6]) was reproduced using the ESMC-6B model (Appendix E). The PPI3D [10] dataset was downloaded with two different date cutoffs: up to 2026-07-29 for exact reproduction of the original method, and up to 2023-06-01 for benchmarking on the Flock Leaked and Leakage-Free Sets, both unclustered, and clustered using 40% sequence identity and 50% interface contact similarity. Normalisation of feature activation signals used the log10 max_idf data. Query pairs were scored as described [6]. For pairs where either chain had no active SAE features, a zero score was imputed. Further detail on the ESMC method and our reproduction of it is given in Appendix E.

Leakage-free pairs above 1,254 residues in size (target + partner combined) were excluded from the benchmark due to available GPU memory.

## 4 Experiments

### 4.1 Creation of Flock, Leaked and Leakage-Free Set

Both Flock and Flock’s Leakage-Free Set contain vastly more negative pairs than previously described datasets (see Table 1). The majority of these negative pairs are retrieved from the PDB (368,937 out of 377,643). Despite the smaller number (8,706), the annotations from literature are confident and high quality, as demonstrated by the lack of conflict with positive filters (PINDER, PPI3D and IntAct). More details of this overlap are described in Appendix D.

**Table 1:** Summary of the number of targets and annotated pairs in Flock and Flock’s Leakage-Free Set compared to previously published benchmarks Negatome v2 [2], PRING [14] and PINDER [9].

| Dataset | No. targets | No. positive pairs | No. negative pairs | Total pairs |
| --- | --- | --- | --- | --- |
| Negatome v2 | 3,217 | 0 | 5,533 | 5,533 |
| PRING | 21,484 | 186,818 | 0 | 186,818 |
| PINDER | 25,269 | **2,319,564** | 0 | **2,319,564** |
| Flock | **34,959** | 26,934 | **377,643** | 404,577 |
| Flock Leakage-Free | 335 | 783 | 11,203 | 11,986 |
| Flock Leaked | 298 | 974 | 12,484 | 13,458 |

The number of positive pairs is smaller than other datasets [9, 14], but this is a result of being derived from experimentally-solved protein structures, as opposed to other indirect evidence of a PPI. Crucially, this allows us to split Flock’s Leakage-Free Set based on interface similarity. This structure-aware split allows the fair evaluation of co-folding methods on the PPI prediction task, particularly for novel, unseen PPI pairs.

### 4.2 Performance of co-folding models and ESMC on Flock’s Leaked and Leakage-Free Sets

**Figure 1:**
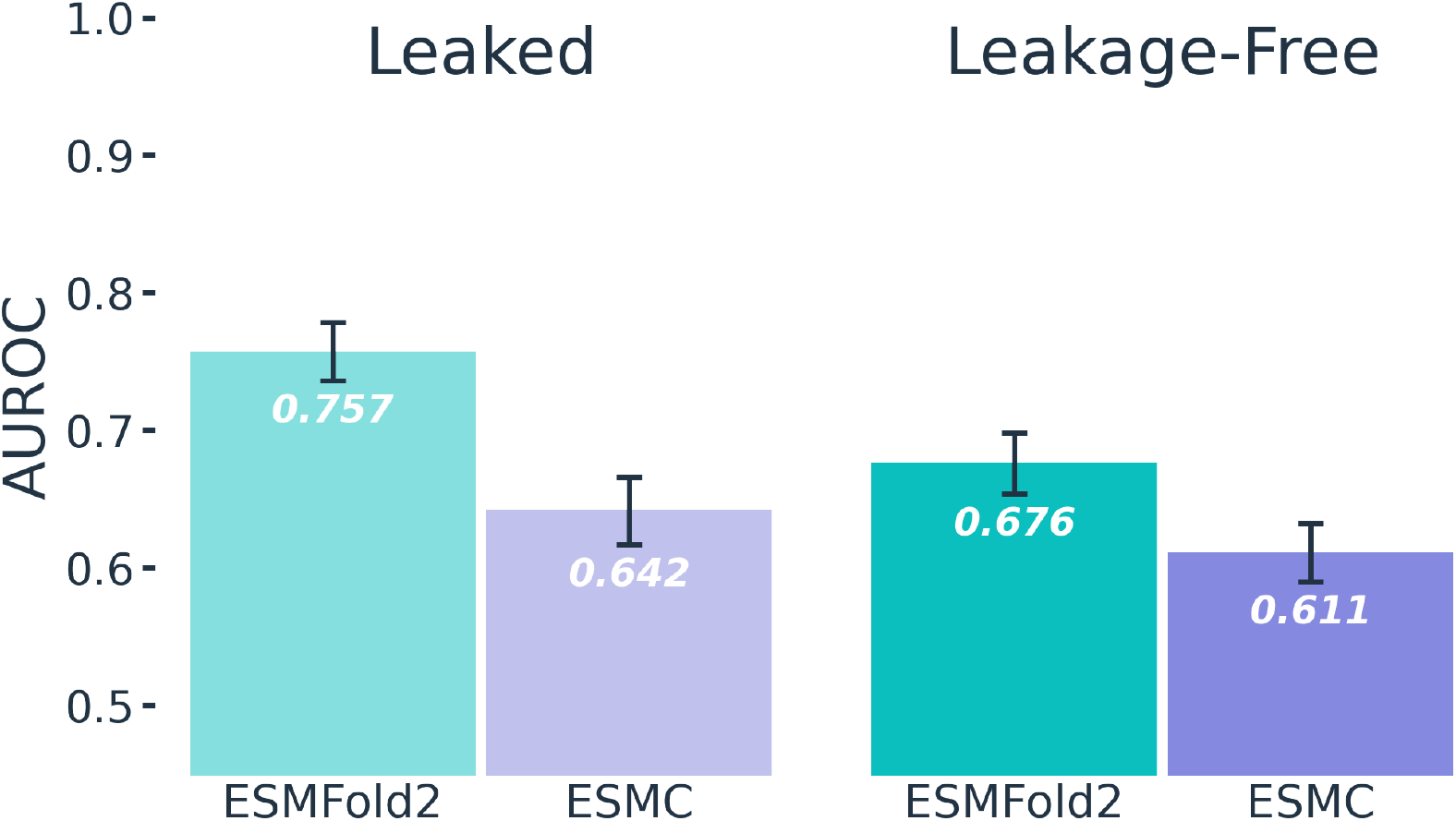
ESMFold2 and ESMC results on Flock’s Leaked and Leakage-Free Sets, as measured by AUROC (values shown as white, italic text). 95% confidence intervals shown as black bars.

Validation of our ESMC reproduction confirmed results comparable to the original publication [6] (detailed in Appendix E). To enable direct comparison with ESMFold2, we applied Flock’s Leakage-Free cutoff date (2023-06-01).

Among all ESMFold2 metrics evaluated on Flock’s Leakage-Free Set by AUROC, the recalculated interface pTM score (`iptm_d0chn`) [17] achieved the highest performance. It outperformed both interface ipSAE (across all PAE cutoffs) and non-interface metrics (PTM and average pLDDT). Additionally, ESMFold2 showed slightly higher performance (Spearman’s *ρ* = 0.195, *p* = 0.001) on targets with higher residue confidence (median pLDDT *>* 65).

Evaluations on Flock’s Leakage-Free Set yielded lower performance for both models relative to Flock’s Leaked Set. ESMFold2 showed a pronounced reduction in performance when evaluating unseen interfaces compared to previously seen proteins (*p* = 2 × 10^*−*12^). In contrast, ESMC had a smaller performance difference between Leaked and Leakage-Free Sets (*p* = 0.027), indicating stronger generalisation capabilities in this protein language model than in co-folding models.

Furthermore, both models showed a substantial reduction in accuracy on the Leakage-Free Set compared to the original benchmarks reported for ESMFold2 and ESMC [6, 18], AUROC dropping from 0.828 for ESMFold2 and 0.855 for ESMC. This performance decrease underscores that Flock’s Leakage-Free Set provides a more rigorous and realistic PPI prediction benchmark by incorporating explicitly defined negative interactions rather than non-positive random pair sampling [18]. This, in combination with consideration of unseen interfaces, gives Flock’s Leakage-Free Set a clear advantage for benchmarking co-folding methods against alternative approaches compared to previous benchmarks.

## 5 Conclusions

In Flock, there now exists a benchmark combining positive and negative (non-interacting) PPIs. The number of negative PPIs identified has increased by 68-fold over Negatome v2 [2]. Using agentic LLM workflows as opposed to semantic natural language processing tools has increased both the number and quality of negative pairs. Flock’s Leakage-Free Set enables fair evaluation of the generalisability of co-folding methods for predicting PPIs. This work demonstrates that failing to account for data leakage leads to overinflated performance for both protein language models and particularly co-folding models.

No negative PPI pair can be established with full certainty, which is an inherent limitation of this dataset. Flock negatives could be as-yet-undetermined positives; interaction between proteins can be dependent on cell type, post-translational modifications, or biological states like disease or stress. This is a limitation of negative interaction data in general rather than of this dataset in particular. Flock’s literature negatives pass through layers of deterministic and agentic curation, and agreement between the literature-mined and PDB-derived negatives gives an independent check on both. The result is a more confident set than that presented in Negatome v2, whose literature-derived pairs include many spurious examples (Appendix A).

Flock’s Leakage-Free Set was used to evaluate two different methods in this work, both of which are based on protein language model embeddings. ESMFold2 trains a co-folding model on these embeddings using protein structures, whereas the ESMC method evaluated here uses a trained sparse autoencoder (SAE) model on the embeddings to learn interpretable feature activations. Of the two methods, ESMFold2 shows a bigger performance drop from leaked examples to Leakage-Free; this is expected since its parameters are directly fitted to this leaked set, while the ESMC method is not trained and only uses it to derive three hyperparameters.

Despite this gap, ESMFold2 still shows better performance on the Leakage-Free Set than ESMC despite using the same underlying embeddings. There are several potential reasons for this: ESMFold2 is trained on protein structures which may encode more meaningful signal on PPIs than sequence alone, or perhaps there are limitations with ESMC’s SAE model used in this method. The original ESMC method chose hyperparameters that were near the edges of the sweep range tested, indicating that these could be further optimised. Finally, for PPI prediction tasks the use of SAE features from protein language models is less well explored in the literature compared to structure-based co-folding models, whose inductive biases and confidence metrics are more well understood.

To conclude, Flock and its Leakage-Free subset represent valuable resources for model training, and challenging, fairer PPI benchmarking. There are many opportunities to develop better PPI prediction models (and better understand their performance) through the use of high quality negative data, diverse literature negatives, and the avoidance of data leakage.

## 6 Data Availability

Code is available from https://github.com/TernaryTx/flock. The Flock dataset is available from https://huggingface.co/datasets/ternarytx/flock.

## A Analysis of Literature Negatives from Negatome v2

We found that the published Negatome v2 dataset from Blohm et al. [2] contains errors and incorrectly labelled negative PPI pairs, a summary of which are shown in Table 2. These errors fall into three groups. Most are mapping failures, where the recorded accession is not the protein the paper discusses: a gene symbol colliding with an unrelated protein, an acronym naming a complex rather than a protein, an affinity tag or reporter carried over from the pull-down, a viral polyprotein standing in for the mature peptide actually assayed, or a splice variant recorded against the canonical sequence. A second group over-generalises, expanding one statement about a protein family into a separate pair for every member. The third inverts the label outright, recording as non-interacting a pair the cited paper shows does interact.

**Table 2:**
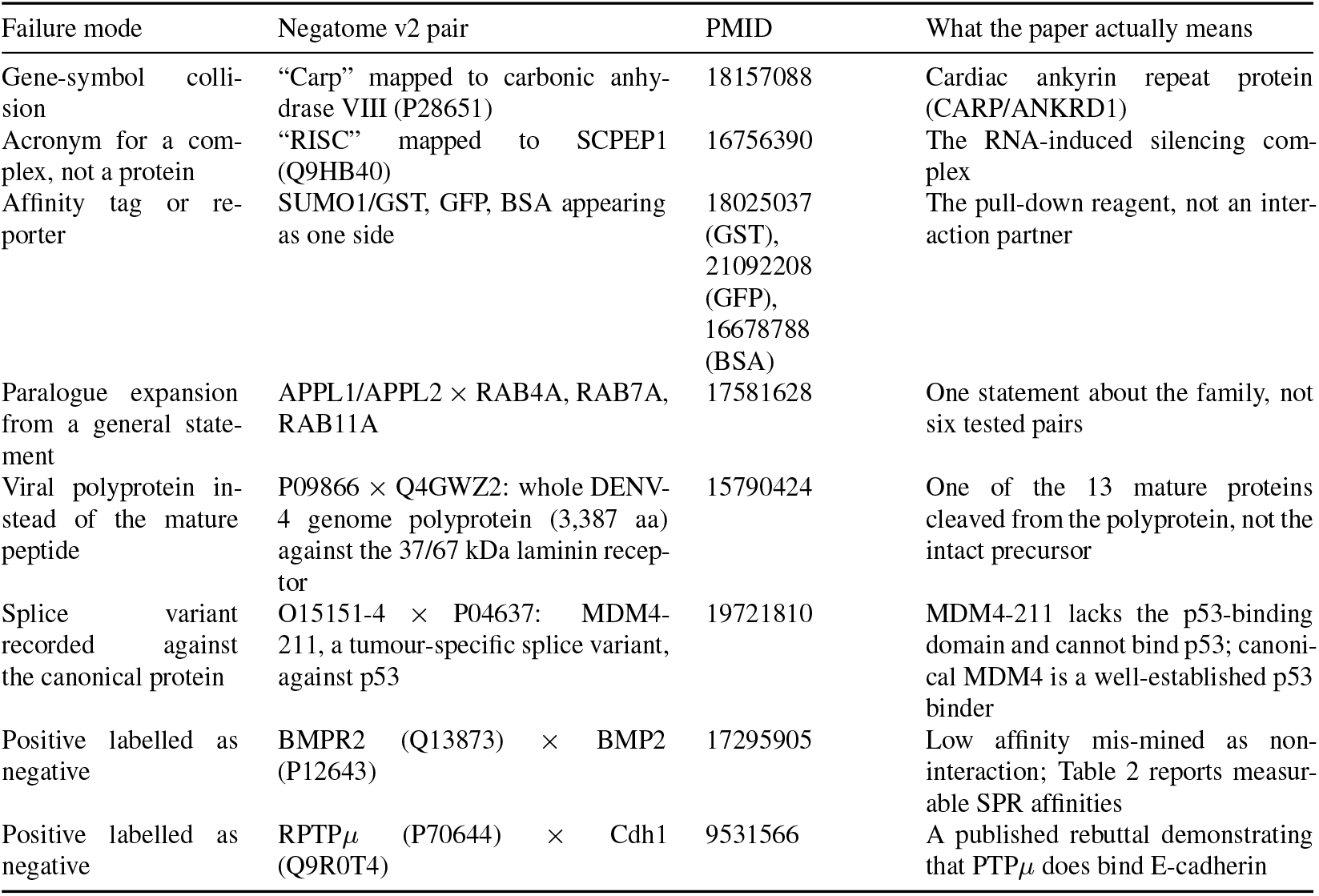
Spurious negative pairs from the existing Negatome v2 dataset. PubMed ID (PMID) is provided for the paper the Negatome v2 based the spurious pair on.

Flock’s literature pipeline addresses each in turn. Names are resolved against an index of reviewed UniProt entries, and names with more than one plausible accession are assigned by a model that sees the paper’s own text. Constructs are filtered before resolution: leading affinity tags and point-mutation notation are stripped, and bare reagent names are excluded. Each accession carries a viral flag, and pairs with no host protein are dropped. Every surviving pair is then judged against the paper that produced it, and admitted only where both accessions match the proteins discussed, the negation concerns the full-length pair rather than a mutant, fragment or family member, and the paper reports the result itself rather than citing it. Finally, pairs recorded as positive in IntAct, or supported by a structure in our positive sources, are removed.

The examples above were found by hand inspection and are illustrative rather than exhaustive; we did not measure an error rate over the published set.

## B Detailed Methods for Retrieval of Literature Non-Interacting Proteins using Agentic Systems

In the Negatome v2 work Blohm et al. [2] applied Excerbt [19], a semantic parsing tool, over PMC abstracts and full texts. They filter to only include negated predicate-argument structures, with proteins in both agent and theme roles, narrowed to interaction verbs (“to interact”, “to bind”, “to co-immunoprecipitate” etc.). Our work differs from theirs for two reasons. Firstly, at the beginning of the pipeline, Europe PMC offers exact-phrase search rather than parsing. Secondly, we eschew tools like Excerbt in favour of LLMs, which can actually reason over text instead of just parsing.

To this end, we search papers in two different ways. We first use a set of exact search phrases to query Europe PMC and decide which papers enter our corpus. For each search phrase, we use a matched regular expression (regex) against the retrieved full text, to decide which specific sentence or paragraph from a paper contains the queried negation phrase/sentence. Every one of the 31 search phrases is matched by at least one regex. Phrases must be exact and adjacent; regexes have slots and tolerate up to three intervening words. Appendix C.4 describes how we arrived at the final set of search strings used.

### B.1 Screening for Negative Pairs using LLMs and Agentic Systems

#### Initial Screen

Claude Sonnet 5 was used through the Anthropic Batch API, with low effort and adaptive thinking, and 16k max tokens for thinking + output. 5,730 papers were refused due to Sonnet 5’s biosafety guardrails; for these examples a fall back to Sonnet 4.6 and (if necessary) Haiku 4.5 was used. Models were constrained to a structured output scheme in JSON containing: a list of pairs with protein names as written in the text, a “relationship” (no interaction, interaction, or both), a confidence rating (high/medium/low), and at least one verbatim excerpt from the text. Responses were submitted with the screening prompt used as the system message, and the paper text provided as a single user message. Two separate LLM screens were run and the union of all pairs labelled as non-interacting was taken forward.

#### Accessions

Claude Sonnet 5 was used through the Anthropic Batch API, on low effort with adaptive thinking and 4k max tokens for thinking and output. The model was provided with the protein name as written, the matched sentence and the text of the paper paragraph containing it, as well as the list of possible matching candidates from the deterministic pipeline. Each of these candidate lines consisted of UniProt accession, protein name, gene names, and organism. Prompt, protein name, texts, and candidates were all provided as a single user message with no system prompts. The model was instructed to abstain from answering if the paper suggested an organism not present in the deterministic list, or if the text of the paper did not conclusively resolve the UniProt from the list provided. The output schema consisted of: the chosen candidate number (0 for abstain), the verbatim text that decided it (empty on 0), and a one-sentence reason.

Of 38,766 named proteins across 17,313 papers, 25,100 (64.7%) returned an accession, 12,838 (33.1%) abstained, 570 (1.5%) had their evidence span fail the verbatim check and were voided, and 258 (0.7%) were biosafety refusals on both Sonnet 5 and Sonnet 4.6. This resulted in a set of 27,429 pairs of UniProt accessions with 63% of these containing an accession retrieved by an LLM. Abstention rate was between 32–37%, and later curation attributed 91.8% of model-picked sides correct against 77.6% for the deterministic text route.

#### Final Curation

In contrast to the previous two LLM steps, curation used an agentic loop through the Anthropic Messages API with streaming. Claude Sonnet 5 was used at xhigh effort, on adaptive thinking with tool calling and multiple turns allowed (max 24k tokens per turn). Each paper was given its own curation run, with the starting request consisting of the prompt and a JSON package containing: pairs to be judged, protein names as written in the text, assigned UniProt IDs, a summary card for each accession (entry name, protein/gene names, organism, length, review status), how each accession was chosen (deterministic/model picked), the text excerpts from the previous screen, as well as the entire paper text minus the methods section. Papers were truncated at 150k characters (affecting 25 papers total out of 14,947). Curation was run locally with no network access given to the model. Two local tools were provided: `uniprot_lookup` for current entries, and `paper_methods`, which returns the omitted methods section in full.

Each curation verdict came back under a JSON schema: a mapping verdict (correct, wrong paralog, wrong protein, wrong species, fragment or isoform mismatch, affinity tag or reporter, cannot determine), the relationship the paper reports (non-interaction, interaction, both, not discussed, or no evidence), a verbatim evidence sentence and its location, a qualifier (full-length pair, domain fragment only, mutant only, different paralog, cited prior work, not a protein pair, indirect or functional only, unclear), a usability flag, confidence and reasoning. The evidence sentence is checked verbatim against the text sent and examples failing this check are removed. We discard the usability flag from the model and decide that a pair is usable if it matches all of these criteria:

- Mapping verdict is correct (wrong species, wrong paralog, cannot determine and the rest all fail)
- Relationship is non-interaction
- Qualifier is full length (so domain fragments, mutants, a different paralog, cited prior work, non-protein pairs, indirect or functional-only claims, and unclear all fail)
- The evidence sentence is a verbatim span of the text that was sent (body text, plus Methods only if the model called the tool for it)

14 papers were permanently refused on biosafety grounds, costing 18 pairs.

### B.2 Deterministic UniProt Mapping

We attempt to map pairs to UniProt accessions using the UniProt reviewed index, release 2026_02, containing 575,503 accessions, 943,159 distinct names, and 14,898 organisms. Each entry contains a gene symbol and every synonym, the protein name and every parenthetical alternative, and the entry-name stem. We first apply filters to remove mutant-only or domain fragment interactions in our dataset of named protein pairs. We use regexes to remove protein names with construct notation (Δ symbols), stated mutants or truncations, residue ranges, point mutations on a name stem, or leading affinity tags. We remove any names caught in these filters which do not otherwise map to any known names in UniProt reviewed. We also remove any pairs whose screen text excerpts do not verbatim map to the paper’s text. These two filters remove 15,743 (10.7%) and 858 (0.6%) of pairs respectively. We then attempt to map normalized protein names (Greek letters spelled out, non-alphanumerics stripped, slash-joined names read both ways) to UniProt entries directly by name.

For proteins which do not map to any UniProt entries by name, we modify the name as written in three ways, and check if this new processed name matches any UniProt entry. We strip prefix two-letter genus codes (for example “AtRAD16” for *A. thaliana* RAD16), single-letter species prefixes (h/m/r/d/x/z/y/b, as in “hMLH1” for human MLH1), and the yeast `-p` suffix convention. We check that the remainder resolves to a UniProt and that the prefix’s taxa actually contain one of the candidates. For proteins which map to several candidate UniProt IDs, we attempt to find organism evidence from the paper text, in order from strongest to weakest evidence: an abbreviation definition in that paper tying this name to an organism, then the species prefix (which must name an organism the paper itself names), then any organism the paper names anywhere. Following attempted organism mapping, we remove any named proteins with more than 50 candidate UniProt accessions as these are generally pan-species protein names (e.g. “Pnp” maps to 990 accessions) or non-protein names (“NADP” maps to 1,128 UniProt accessions).

Each protein name is then given an outcome. Candidate accessions which differ only by organism share an entry-name stem (for example “SUMO1_HUMAN” and “SUMO1_MOUSE”), and we treat a name resolving to a single stem group as determined rather than ambiguous. Names are therefore labelled as determined, a species pick (the same protein in several organisms), an identity pick (different proteins sharing a name), too wide (more than 50 candidates), or unresolved (no candidates at all). Each pair takes whichever of its two names needs more work. Pairs whose names are both determined are fully mapped, pairs whose harder name is a species or identity pick are passed to the LLM screen described in Appendix B.1, and pairs carrying an unresolved or too wide name are set aside without further mapping.

Finally, we apply two filters which require mapping to have been attempted first. We remove pairs naming an assay reagent or affinity tag (GST, BSA, MBP, DNA and similar), matching against the whole normalised name rather than any substring, since a substring match also removes legitimate entries (for example “NADP-dependent malic enzyme 4” matches “NADP”). Mapping does not substitute for this list, as GST resolves to a real human glutathione S-transferase. We also remove pairs in which both proteins are viral, keeping host-pathogen pairs. These two filters remove 3,025 (2.32%) and 162 (0.12%) of pairs respectively, leaving 127,289 pairs of 130,476: 10,152 fully mapped by name and paper text alone, 31,519 with at least one ambiguous name, and 85,618 unresolved.

## C Literature Screen Agentic System Tuning Experiments

### C.1 Test Corpus

We assembled a working set of 1,155 papers, consisting of all 652 papers cited by the existing literature Negatome v2, and 503 cited by positive interactions from IntAct. Because publication year drives both open-access availability and writing style, IntAct papers were sampled to reproduce the Negatome papers’ year histogram (1975 to 2011 with a median around 2004), and weighted within each year toward papers reporting few interactions each — 405 low, 76 mid, 22 high throughput. We deliberately down-weight high-throughput screens since these papers less commonly mention individual protein–protein pairs.

Full text could be downloaded from Europe PMC for 182 papers from this set (145 Negatome, 37 IntAct). A further 348 were indexed and searchable on their body text but not downloadable, and 624 were absent from Europe PMC. From 182 downloadable papers we hand-annotated 101 Negatome v2 papers drawn at random plus all 25 IntAct control papers in the sample; this gave a set of 262 annotated protein pairs in total.

We use the regex pre-pass to surface candidate negation sentences for the curator to read (Table 3), but 20% of the queue was annotated blind, with the candidate list hidden. This blind subset estimates how often a regex misses the negation statement. The curator annotated each pair with the verbatim sentence claiming the pair, as well as the section of text it came from, whether the pair was positive or negative, whether UniProt mapping may be incorrect, and general confidence (low/medium/high). A summary of the manual annotation results is shown in Table 4.

**Table 3:** Example regular expressions used to identify candidate negation phrases in the literature test corpus. These follow the “negation-plus-interaction-verb” rule described by Blohm et al. [2]. The full set is 15 patterns across the 5 types.

| Phrase type | Example regex (fragments) |
| --- | --- |
| explicit_no_interaction | co-?immunoprecipitate\|co-?precipitate\|<br>co-?localis\|co-?localiz\|complex)\w* |
| failed_assay | \b(?:unable\|inability)\s+to\s+<br>(?:\w+\s+){0,2}?(?:interact\|bind\|associate\|detect)\w* |
| negative_result | \b(?:undetectable\|not\s+detected\|<br>below\s+(?:the\s+)?detection)\b |
| elliptical_exclusion | \b(?:interact\|bind\|bound\|binding\|associat)<br>\w*[\^.] {0,120}? |
| contrast_negative | \b(?:whereas\|while\|but\|however\|<br>in\s+contrast)\b[\^.] {0,80}?\b |

**Table 4:** The number of curated pairs with each type of annotation outcome. Unusable non-interacting pairs include those with incomplete/low quality evidence, evidence cited from other papers rather than shown in the paper itself, or negatives based on domain fragments, mutants, isoforms, or viral polyproteins.

| Annotation outcome | No. of pairs |
| --- | --- |
| Stated non-interaction and usable | 78 |
| Stated non-interaction and unusable | 74 |
| Positive interaction | 63 |
| Pair never discussed in paper | 25 |
| Paper reports both interaction and non-interaction | 12 |
| Positive pair labelled as negative by paper | 6 |
| No/insufficient evidence either way | 4 |

We hand annotated each paper using a Jupyter widget and marked any unsure examples. We then batch reviewed 163 pairs the curator flagged using a Claude Opus 5 agent, starting with unsure examples. Each pair was sent as a self-contained packet with no network access; this packet contained the paper’s metadata, the curator’s own verdict and free-text note, UniProt summary cards for both proteins (sequence length, organism, gene names, cross-references) and the full text of the paper. The agent returned the following information for each pair:

- Whether the UniProt mapping was correct, i.e. one of: correct, wrong protein, wrong paralog, fragment or isoform mismatch, affinity tag or reporter, cannot determine.
- The relationship the paper actually reports: non-interaction, positive interaction, both reported, not discussed, no evidence.
- An evidence sentence and the section it came from.
- What the statement is about: a full length protein pair, domain fragment only, mutant only, different paralog, cited prior work, or unclear.
- Whether the pair is usable as a negative based on the above.
- The agent’s own confidence; whether it agreed with the curator; and its reasoning.

All 147 quoted sentences were machine-checked against the packet they each came from, and none were fabricated. The agent agreed with the curator on 67, partially agreed on 67 and disagreed on 29. Where agent and human agreed, we kept the judgement as is. We manually reviewed any disagreements, most of which were accepted, but a few needed minor tweaks. Of 163 pairs, the agent judged the UniProt mapping correct on 113 and wrong or undeterminable on 50, and 23 survived as usable negatives.

The 126 annotated papers were split into a testing set of 83 papers and a held-out set of 43. Within the tuning set, 52 annotated pairs were designed as pairs a screen ought to surface (41 stated non-interactions and 11 pairs the paper reports both ways). 49 pairs were those that an LLM must not surface as negatives — positive interactions from papers that nonetheless contain negation language. We use this set of 83 papers and 52 desired pairs/49 undesired pairs to run the experiments in the next sections.

### C.2 Experiment 1: Model Choice

As shown in Table 5, we tested Haiku 4.5 vs Sonnet 5 with low effort or no thinking enabled. We found that no thinking performed worse than low effort — it recovered the same 37 pairs as the low setting while generating 62% more tokens, quoting absent evidence three times as often (6 against 2), and returning 45% more unlabelled pairs (824 against 570). Since downstream curation is also a cost (in terms of tokens and human effort), generating more pairs for the same recall is more expensive. Both Sonnet 5 settings refused the same paper on safety grounds — a study of the HIV envelope trimer (1 of 83). Haiku 4.5 refused no papers.

**Table 5:** Results of using different Anthropic models on literature test sets.

| Model | Pairs found (of 52) | Incorrectly found (of 49) | Evidence not in text | Unlabelled pairs |
| --- | --- | --- | --- | --- |
| Haiku 4.5 | 42% (22) | 0% (0) | 11 | 654 |
| Sonnet 5, no thinking | 71% (37) | 0% (0) | 6 | 824 |
| Sonnet 5, low effort | 71% (37) | 2% (1) | 2 | 570 |

Note that repeat runs of one configuration do not return identical results. Three replicates with Sonnet 5 low effort on the same prompt and input recovered 39, 41 and 44 of the 52 pairs. 39 were found by all three and 5 by some runs but not others. Sonnet 5 does not accept a temperature setting, so this variation can be measured but not suppressed. The 15-pair gap between Haiku 4.5 and Sonnet 5 clearly separates from this background variation, however.

### C.3 Experiment 2: Model Inputs

In this work, we search papers in Europe PMC for negation phrases in sentences, before using matched regular expressions to extract those specific sentences or phrases. It is therefore possible to extract a matched paragraph for any negation statement found, meaning any paragraph containing the negation expression. This could save input tokens but may also affect model performance. We compared whole papers (minus methods sections) against only matched paragraphs to investigate where negation claims are made and how this affects model performance.

From the hand annotated set only 11% of non-interaction claims are stated in the abstract, rising to 83% once the body is included and 88% with figure and table captions; the remaining 12% are in supplementary material or otherwise unrecoverable. Targeting named sections rather than the whole body caps at 88%, because around 20% of articles use topic-based section headings that cannot be classified as Results or Discussion at all. An independent check from search counts agrees: “does not interact” returns 30,936 papers, but only 2,535 when restricted to abstracts (8.2%).

We then checked whether each annotated negation phrase/sentence could be recovered from matched paragraphs or sentences. As shown in Table 6, we found that extracting just the matched paragraph surrounding a regex provides the best balance between performance for the amount of text used. We briefly experimented with extracting candidate sentence spans but this was found to degrade performance too much (data not shown).

**Table 6:** Recovery of negation sentences with different amounts of text used.

| Text sent | Mean characters | % of whole paper (no Methods) | Pairs evidenced (of 90) |
| --- | --- | --- | --- |
| Whole paper (no Methods) | 35,400 | 100% | 99% (89) |
| Matched paragraphs ($\pm 1$) | 15,100 | 43% | 96% (86) |
| **Matched paragraph only** | **7,500** | **21%** | **91% (82)** |

We then investigated how these different text splits affected model performance. Sonnet 5 on low effort, given whole papers (no methods), surfaced 37 of 52 annotated test pairs. The same setup with matched paragraphs surfaced 41 pairs (≈10% gain) while sending only 21% of the text. The total number of prospective pairs emitted fell 17% and pairs quoting absent evidence fell from 2 to 0, so the reduction also cuts the downstream review workload. This performance gain is within variation as seen in the previous section, but notably results in lower projected cost, with full text averaging around 13,545 input and 1,142 output tokens per paper to matched paragraph’s 3,417 and 925.

We carried out a sweep across effort levels for this step, and found that low, medium and high effort all recovered the same pairs across a 3.4-fold range of output tokens. Only xhigh did better, by two pairs, but at an estimated 6× cost increase. On low effort, a second screening prompt, tested with the output schema held fixed, recovered 44 of 52 pairs but found exactly the union of what three replicates of the original recovered, so it surfaces nothing the shipped prompt cannot. It also emitted 31% more negative calls per paper, and in one case merged fragment-level evidence into a claim about the full-length pair.

### C.4 Experiment 3: Search Strings

Previous experiments started with a known corpus of 652 Negatome v2 papers. In production, we retrieve our starting corpus by searching Europe PMC with exact phrases, so the quality of these search strings affects our entire dataset. We start with phrases constructed from negation forms (“did not”, “does not”, “was not”, “could not”, “failed to”, “unable to”, “no”, “absence of”, “lack of”) crossed with Negatome v2’s interaction verbs (“interact”, “bind”, “associate”, “co-immunoprecipitate”, “pull down”, “form a complex”, “detect an interaction”), keeping grammatical combinations.

We searched against the entire Europe PMC corpus, as well as the existing 652 Negatome v2 literature papers, to measure both how many known papers a phrase matched, as well as the total number of hits. This allowed us to estimate both the signal and noise returned by a given search string.

The 9 groups shown in Table 7 are exact phrases, matched by exact adjacency. The negation regexes used later to extract a matched paragraph (Table 3) are more permissive — they tolerate up to three words between the negation and the verb. This means that any adverb in that gap (“does not directly interact”) names a real claim the phrase set cannot retrieve, however well the expression matches it once the paper is already in hand. We tested this gap directly (Table 8) with 68 additional candidate phrases. These include adverb insertions, co-precipitate spelling variants, and a few nominal forms.

**Table 7:**
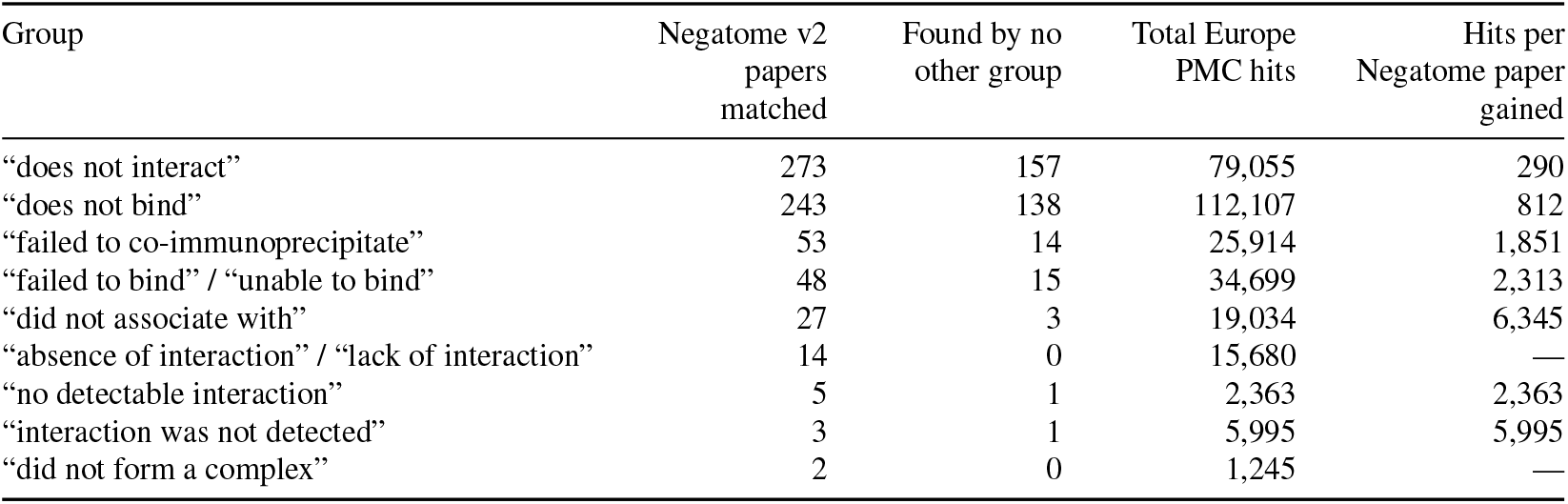
Initial search phrases and their hits against all of Europe PMC as well as Negatome v2 papers. Note that groups of phrases contain more than the one phrase shown here.

| Group | Negatome v2<br>papers<br>matched | Found by no<br>other group | Total Europe<br>PMC hits | Hits per<br>Negatome paper<br>gained |
| --- | --- | --- | --- | --- |
| “does not interact” | 273 | 157 | 79,055 | 290 |
| “does not bind” | 243 | 138 | 112,107 | 812 |
| “failed to co-immunoprecipitate” | 53 | 14 | 25,914 | 1,851 |
| “failed to bind” / “unable to bind” | 48 | 15 | 34,699 | 2,313 |
| “did not associate with” | 27 | 3 | 19,034 | 6,345 |
| “absence of interaction” / “lack of interaction” | 14 | 0 | 15,680 | — |
| “no detectable interaction” | 5 | 1 | 2,363 | 2,363 |
| “interaction was not detected” | 3 | 1 | 5,995 | 5,995 |
| “did not form a complex” | 2 | 0 | 1,245 | — |

**Table 8:**
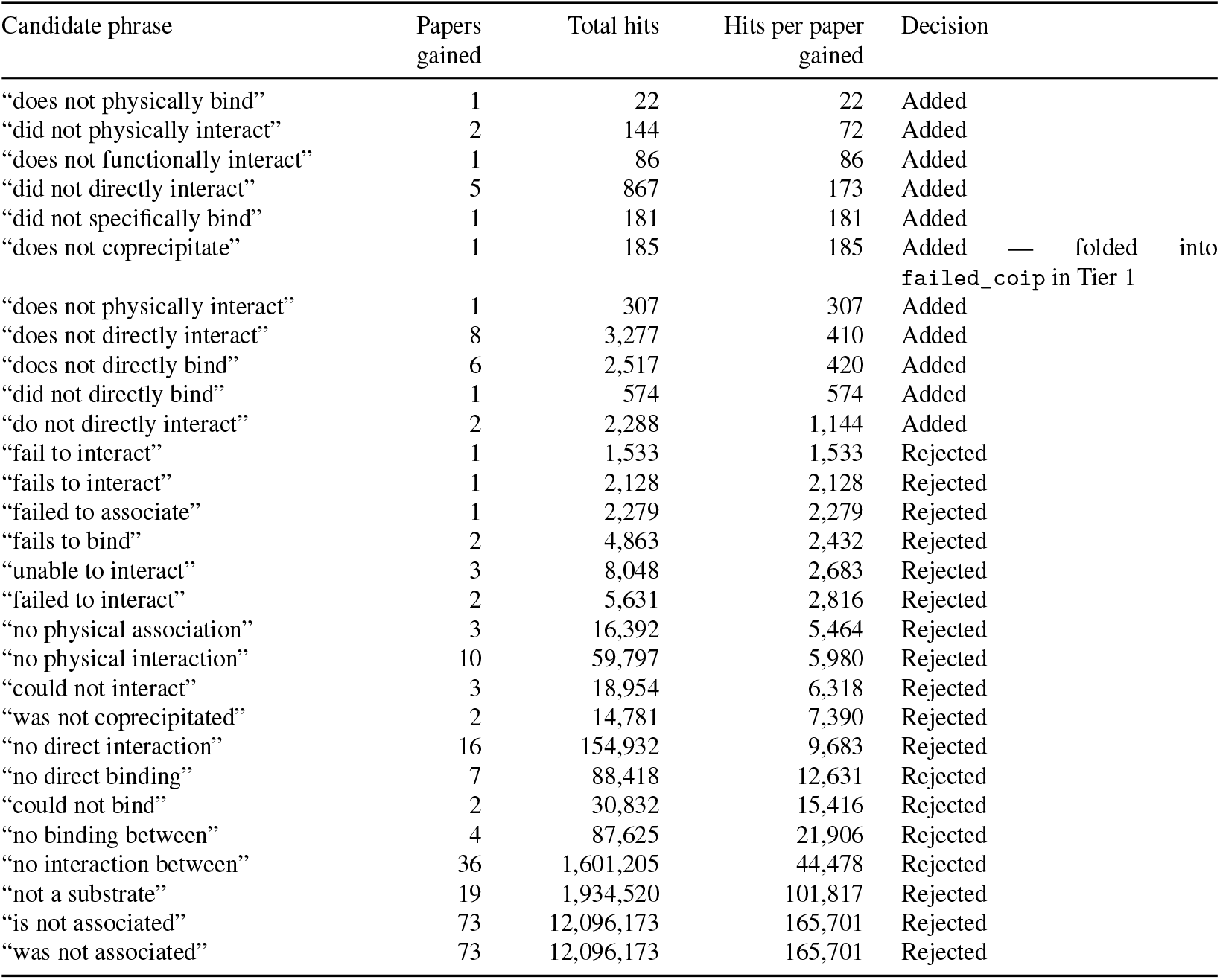
Additional candidates tested against papers not surfaced by the 9 groups in Table 7. Not shown: a further 39 candidates which surfaced no previously-missed papers (hit counts 2 to 19,197). These 39 candidates were mostly systematic inflections of the added phrases (“do”/”does”/”did” + the same verbs) plus a few nominal forms (“cannot bind”, “no evidence of interaction”).

These new candidates were screened against the papers the 9 initial groups did not already match, again scoring each on papers gained and total hits. The missed papers were split by identifier hash and only one set was read when mining for new phrasings. Of the 29 new papers, 15 recovered papers fell in the unread set, and a further 4 came from phrases generated by the systematic adverb grid alone, and never observed in any read text. As such these additions generalise rather than fitting to the 172 papers we mined them from. We decided to keep new candidate phrases whose cost (in terms of hits per known paper surfaced) fell in line with the existing seed phrase set.

Finally, we pruned some phrases from the screening set based on leading negative terms. Europe PMC’s phrase index discards a leading “no”, “not” or “a”, so a phrase that begins with these negations retrieves exactly the same papers as the phrase with the negation removed. This is position dependent: “does not interact” gives 30,936 hits vs 2,022 from “does interact”, but conversely “no detectable interaction” and “detectable interaction” both return 2,428. Two phrases in Table 7 fail this test (“no detectable interaction” and “was not co-immunoprecipitated”) but were kept anyway for the 5 and 51 known papers they respectively reach. The 9 phrases in Table 9 either failed this test, or failed to retrieve any additional papers not recovered by remaining phrases, and were dropped.

**Table 9:** Phrases dropped from the original seed set. Seven of the nine fail to select on negation (Europe PMC returns the same papers with the negation stripped); the other two are selective but contribute no known paper beyond the shipped 9-group set. Dropping all nine costs 4 of the 652 known Negatome v2 papers, all from phrases that also fail the negation test.

| Phrase | Negatome papers matched | Unique papers added | Total hits | Hits with negation removed | Reason for removal |
| --- | --- | --- | --- | --- | --- |
| “was not pulled down” | 45 | 2 | 44,961 | 44,961 | No different with negation removed |
| “did not pull down” | 3 | 0 | 1,351 | 81 | Adds no new papers |
| “failed to pull down” | 1 | 0 | 865 | 865 | Adds no new papers |
| “not a direct interactor” | 1 | 0 | 630 | 630 | No different with negation removed |
| “not an interacting partner” | 9 | 2 | 12,716 | 12,716 | No different with negation removed |
| “no detectable binding” | 10 | 0 | 7,760 | 7,760 | No different with negation removed |
| “no significant binding” | 8 | 0 | 17,673 | 17,673 | No different with negation removed |
| “no detectable association” | 0 | 0 | 1,305 | 1,305 | Matches no known paper at all |
| “no interaction was detected” | 11 | 0 | 12,963 | 12,963 | No different with negation removed |

### C.5 Experiment 4: Abstract vs Full Text Papers

Europe PMC hosts a mixture of open access as well as paywalled articles. The body text of all papers is indexed and searchable, but the full text is only downloadable for open access articles. For example, the search phrase “failed to co-immunoprecipitate” matched 448 papers, of which only 200 are open access. For the rest, only an abstract is available. Consider a paywalled paper with a sentence claiming that two proteins do not interact, found in the results section. Our search string would only return the abstract of this paper, which may or may not restate this non-interaction.

To avoid including every paywalled paper we retrieve and checking whether the claim survives into the abstract, we investigate whether we could split retrieval into two routes to potentially save on model screening later. For open-access papers we fetch full text and screen the matched paragraph, as described above. For paywalled papers, we restrict the search itself to the abstract field, so that any match retrieved is provably present in the abstract text itself.

The value of this split is shown on the known Negatome v2 paper set. From 504 known papers our search strings retrieve, only 137 are open access. Abstract-only search reaches a further 301, leaving just 66 papers whose only match is found in paywalled text. To quantify the performance and cost of this abstract-only search, we randomly sample 1000 papers from our search corpus — 600 open access papers and 400 paywalled papers, published 2010 onwards, and matched across four publication-year bins.

Table 10 shows the results of this experiment. While abstract-only search returns far fewer candidate pairs per paper, its projected cost is only ≈11% of the bulk open access screen. We therefore decided to include the abstract-only screen in addition to the open access route. In this experiment we observed that yield is heavily concentrated. On the open-access route, 301 of 499 answered papers yielded no negative at all; the median is zero, the 90th percentile 4, the 99th 9, and one paper produced 22. Papers yielding five or more account for 47% of all negatives found.

**Table 10:**
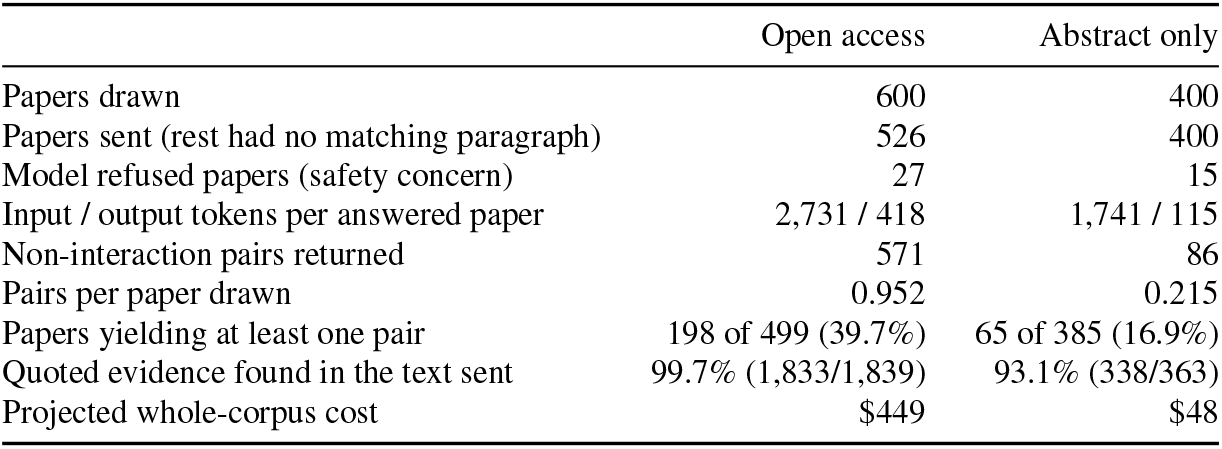
Performance and cost of open-access and abstract-only screening, measured on 600 open-access and 400 abstract-only papers drawn from the production corpus (published 2010 onwards, matched across four publication-year bins so neither route’s numbers are confounded by publication era). Abstract-only screening costs about 11% of the open-access route’s projected bill while returning roughly a quarter as many pairs per paper drawn (0.215 vs 0.952).

Abstract-only screening and the experiment presented here come with several limitations. Firstly, we cannot estimate the recall of actual, real negative pairs by this method, since none of the 78 usable negatives in our annotated set is stated in an abstract. Since the sampled papers in this experiment post-date our Negatome v2 annotated set, we cannot measure precision directly either, and instead use proxy metrics — quoted-evidence checks and yield per paper. In our annotated set, no abstracts establish the three commonest reasons a stated non-interaction is unusable: that it concerns a truncated construct, the wrong paralog, or an interaction mediated by a third protein.

Finally, we observe substantial run-to-run variation in this experiment. We note that on two separate runs of the abstract-only search, 86 non-interacting pairs were returned from the same 385 papers, but only 63 of the 109 pairs in their union appear in both. 42% of the union was found exactly once; the Jaccard coefficient is 0.578, and restricting to the model’s own high-confidence calls does not improve it (Jaccard coefficient: 0.568). Compared on pair counts alone the two runs would have appeared perfectly reproducible. Refusals, by contrast, are near-deterministic: 14 of 15 shared. A single screening run is one sample of the negatives in a corpus, not the negatives in that corpus.

As such, we decided to include abstract-only screening since it is very cheap and appears to perform well enough, albeit as measured indirectly. However we acknowledge that it may introduce some low-quality pairs into our downstream set which require more curation.

### C.6 Experiment 5: UniProt Accession Mapping

Following the initial LLM screen of our paper corpus, we retrieve pairs of candidate non-interacting proteins, as named in the text. We attempt to deterministically map these named proteins to a single UniProt accession, but in many cases there are multiple matching UniProt candidates for a given protein name. For example, the name “Cdc42” in text matches different UniProt entries including Q05062 (*C. elegans*), P60953 (human), P19073 (*S. cerevisiae*) and so on. Similarly the name “CRP” matches human C-reactive protein (P02741) or *E. coli* catabolite activator protein (P0ACJ8). We attempt to resolve the former case by searching for species text in the paper but this is not always possible, and we have no way of deterministically solving the latter problem. As such we attempt to solve both problems using an LLM to assign UniProt accessions from a list of matching candidates.

To determine the best model and inputs for this task, we ran an experiment on 152 pairs from 74 papers in the curated set, where for every named protein we have a curator-confirmed UniProt accession, but multiple candidates emitted from our deterministic pipeline on the protein’s name as written in the text. We test Claude Sonnet 5 and Haiku 4.5, providing each model with either the extracted sentence from the screen only, the paragraph containing this sentence, or the full-text paper. We cap thinking and output tokens at 4,000 and use adaptive thinking and low effort. The output schema was limited to three fields: the chosen UniProt accession, the verbatim sentence from the text that explains it, and a one-sentence reasoning for the choice. Models were given the choice to abstain. The results from this experiment are shown in Table 11.

**Table 11:**
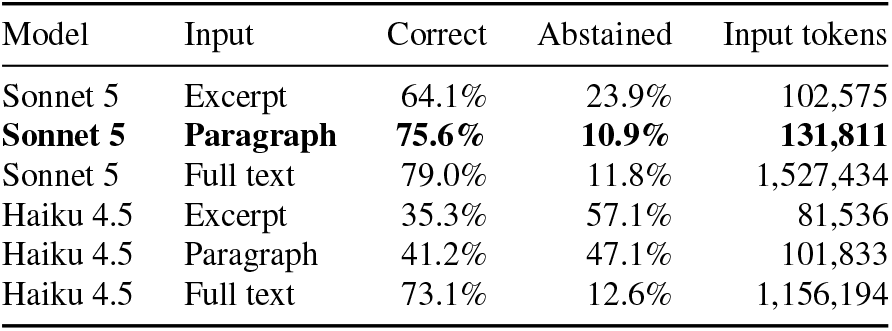
Experiments using LLMs to assign UniProt accessions to curated pairs (152 rows from 74 papers of the curated set). Bold indicates the scheme chosen for the production run.

We find that Haiku 4.5 shows poor performance on this task, abstaining and refusing to answer entirely in many cases. Performance was better when provided with full text, but at ≈10× cost in terms of input tokens. By contrast, Sonnet 5 showed much better performance with all three input levels. As Table 11 shows, we chose paragraph inputs over full text since it offers comparable performance, but with the aforementioned ≈10× saving of input tokens. However we note that there is a degree of non-determinism in this setup, as is expected from LLM pipelines. We ran the chosen Sonnet 5 experiment in duplicate and, whilst the correct rate was exactly the same (76.7%), 13 rows gave a different answer in both runs. 10 rows were correct in one run but not both, however 10/14 of the incorrect rows were incorrect in both runs, meaning the model generally repeats mistakes. We investigated only accepting UniProt picks where both runs agreed, but found that the correct rate increased by 3.0 percentage points, while the answer rate fell 7.8 points. We also investigated asking models to return a confidence field, but found that it failed to distinguish between correct and incorrect choices on a held out set of 27 rows, and increased the abstention rate from 11.2% to 19.3% with no gain in accuracy.

### C.7 Experiment 6: Final Curation of Negative Literature Pairs

Finally, we investigated the use of agentic systems to curate our dataset of putative negative protein pairs, using the setup prototyped in Appendix C.1 as a starting point. We took a sample of 40 papers containing 127 pairs from our production LLM screen, stratified by paper length and by how many pairs each paper carries, with the length tertiles cut on all 21,589 candidate papers. Four abstract-only papers were included. Note that this dataset has no confirmed ground-truth labels, and is used to measure disagreement between methods and cost, not correctness or accuracy. All experiments used Sonnet 5.

We first prototyped this stage using headless Claude Code invocations (claude -p) at $0.67–$1.00 per paper. We found that most of this cost was from input tokens from the harness and system prompt (one run billed over 100k input tokens for an 11k token packet). The same experiment run through the Messages API cost around $0.05 per paper. We test a single-pass curation run against agentic multi-turn loops as shown in Table 12. We find that a single-pass run generates the most negatives labelled as usable at the lowest cost, and that agentic curation (regardless of effort) is consistently stricter. Agentic xhigh outputs 16% fewer labelled negative pairs, mostly reclassifying a claim as cited from another paper or as concerning a domain fragment where cheaper settings called it a full-length pair. We also note that low effort models do not use tools, even in the agentic run. This is because, as described in Appendix C.1, we provide the agent with a UniProt data packet for the chosen pair. It is only on xhigh effort that the model reasons enough to make external tool calls and check the validity of this context. We also observed in the single-pass run that 2/40 papers returned 0 output tokens and no verdict, with no error raised, and were billed for input tokens. As such, we check the stop_reason field returned by the API in production and retry empty papers.

**Table 12:** Curation experiments over 40 papers and 127 pairs. Tool calls are totalled across all 40 papers. Projected cost is calculated over the estimated 15,744 papers from the full production run. Only single-pass curation can make use of the Batch API at 50% discount, since multi-turn loops cannot be batched. Bold indicates the scheme chosen for production.

| Configuration | \$/paper | Output tokens per paper | Turns | Tool calls | Projected corpus cost | Usable negatives kept | Verdicts differing from single-pass low |
| --- | --- | --- | --- | --- | --- | --- | --- |
| Single pass, low effort | \$0.050 | 1,003 | 1.00 | 0 | \$395 | 70 | — |
| Agentic, low effort | \$0.052 | 1,138 | 1.03 | 0 | \$819 | 66 | 12% |
| **Agentic, xhigh effort** | **\$0.109** | **5,773** | **1.78** | **34** | **\$1,718** | **59** | **28%** |

## D Overlap Between Identified Negatives and Positive Datasets

Negative pairs that contradict with positive pair data were removed. Table 13 shows the number of pairs removed using each of the positive source filters broken down by negative source. It can be seen that no literature pairs are found in the positive source data.

**Table 13:** Overlap between positive sources of data (IntAct, PINDER and PPI3D) vs Flock’s negative sources of data (PDB and the literature — Lit). Each row is the number of pairs removed sequentially in the pipeline (e.g. 22,403 pairs removed first by the IntAct filter, then 1,009 pairs are removed using the PINDER filter of the remaining pairs).

| Positive source | Negative source |  |
| --- | --- | --- |
|  | PDB | Lit |
| IntAct | 22,403 | 0 |
| PINDER | 1,009 | 0 |
| PPI3D | 225 | 0 |
| Total | 23,637 | 0 |

## E Details of the ESMC Method and Our Reproduction

### E.1 Model weights, SAE and feature extraction

ESMC was reproduced broadly following the methodology described [6]. We utilised the `biohub/ESMC-6B` model weights and `biohub/ESMC-6B-layer60-k64-codebook16384` sparse autoencoders (SAE), as the normalisation statistics (IDF, max) for these data are publicly available at https://biohub.ai/models/esmc. We maintain the activation threshold of 0.5, storing the top-*k* number of features set to *k* = 64 as determined by the SAE from ESMC. At the point of extraction for query pairs (as a lead time truncation), we store the top 15 features, before scoring on the optimised parameter from the ESMC paper of `top_k` = 3. Every run is fingerprinted and checked before scoring, which ensures that a score can only be looked up in a table built from the same feature space.

### E.2 Normalisation statistics

For normalisation, we use the log10 `max_idf` statistics (md5 `b75317aab86ba3eecd696b8d8ff2ba2e`). This is due to closer reproduction of the paper’s “∼1 million” feature pairs in the NPMI table. Use of the natural log files provided gives a table roughly 8 times the size, with a less even distribution of metrics resulting in many being close to the threshold for active or not. Using log10 metrics provides a table of 631,509 feature pairs, which was hence taken forward.

### E.3 PPI3D corpus

For the reproduction of the ESMC paper [6], we download an unclustered dataset from PPI3D [10], using a cutoff of 2026-07-29. The exact date is not stated in the original paper for the cutoff. This gives us the fullest PDB data available at the time of this study for analysis. For benchmarking purposes against Flock’s Leakage-Free and Leaked Sets, we use the cutoff of 2023-06-01 for consistency.

Prior to clustering, protein–nucleic acid complexes are removed from the dataset, leaving us with protein–protein and protein–peptide complexes. This removal must be done prior to clustering since “cluster type 1” holds both the peptide and nucleic acid complexes together. Consequently, filtering after clustering may delete peptide complexes that are included in a nucleic acid representative.

The PPI3D corpus is clustered at 40% sequence identity and 50% interface contacts using PPI3D’s clustering label of `“cluster_data_40”`. Where needed, representatives from each cluster are selected first by best resolution, and then largest interface area, if there are any exact matches. The resulting sizes of the corpora are as follows: 2026-07-29, 631,509 feature pairs; 2023-06-01 strict cutoff (with removal of any clusters that would have contained any PPIs from after the cutoff), 478,830 feature pairs. Interface contacts are scraped from the PPI3D per-interface details pages keyed by internal database IDs, and subsequently joined after SAE feature extraction. NPMI lookup tables are generated following the equations detailed in the ESMC paper [6].

### E.4 Scoring

The hyperparameters for all experiments in this study are fixed to those described as optimal in the ESMC paper [6]. These are as follows: `top_k` = 3, *ρ* = 0.25 and *T*_max_ = 70. Other parameters fixed in equations are also maintained for consistency (*α* = 0.1, `min_npmi` = 0). Given that the scorer returns NaN when either protein has no residues with active features, we imputed zero as a conservative choice. There are more genuine NPMI = 0.0 scores in the dataset given none of the selected feature pairs carried a positive NPMI; these ones are not imputed. We calculate AUROC measurements for all test sets with and without zeros imputed to ensure that the data are not drastically skewed by this imputation. Query scoring code is provided alongside this study.

### E.5 Evaluation on the Zhang et al. [18] test set

The benchmark used in the ESMC paper [6] came from the Zhang et al. [18] study (their Supplementary Table 1). The test set contains 3,000 positive PPIs and 30,000 random negatives. Using MMseqs2 [20] (30% sequence identity, 80% coverage, non-inclusive, –`dbtype` and –`search-type` set explicitly to amino acid), the pairs are filtered. The ESMC paper retained 778 positive pairs and 12,355 negative pairs. In our reproduction (and due to differences in the overall corpus), we retain 738 positive pairs and 11,897 negative pairs. Based on these data, we report an AUROC of 0.902 (zeros imputed) and an AUPRC of 0.542. The ESMC paper reported an AUROC of 0.855 and an AUPRC of 0.467. We believe therefore to have successfully reproduced the PPI scoring method detailed in the ESMC paper.

### E.6 Flock benchmark evaluation

The Flock Leaked and Leakage-Free Sets were prepared as detailed in this paper. For the pre-cutoff strict corpus (ending 2023-06-01), we report AUROC [95% confidence intervals] of 0.611 [0.590, 0.632] on the Leakage-Free Set over 292 targets, and 0.642 [0.617, 0.666] on the Leaked Set over 298 targets.

We also ran ESMC on Flock’s Leaked and Leakage-Free Sets using the full corpus (ending 2026-07-29). In this study we report an AUROC of 0.637 for the Leakage-Free Set, and 0.649 for the Leaked Set. Since the leakage-free/leaked split is defined against the co-folding training dates, we do not report further on this study. The similarity in AUROC values may reflect the fact that ESMC had access to sequences in the Leakage-Free Set in this instance.

### E.7 Reproducibility

Data has been made available including code for scoring query pairs, both NPMI look-up tables (full corpus and pre-cutoff) and Zhang et al. [18] test set sequences pre-clustered, alongside the results of our test set and benchmark Flock studies (see Section 6).

Key points for reproducibility:

- For SAE feature extraction, version 3.3.0 of esm must be pulled to align with our NPMI tables. Version 3.4.0 and beyond pulls xformers in as a transitive dependency, which changes the feature space and invalidates the existing NPMI table.
- fp32 versus bf16 moves some (0.44%) of features across the 0.5 activation threshold.
- There is variability in the log10 versus natural log normalisation as noted; we have used log10 normalisation files and this must be used for accurate query searches to be made.

